# Function-first discovery of high affinity monoclonal antibodies using Nanovial-based plasma B cell screening

**DOI:** 10.1101/2024.08.15.608174

**Authors:** Dilip Challa, Joseph de Rutte, Cassady Konu, Shreya Udani, Jesse Liang, Patrick j. Krohl, Ronald Rondon, Kent Bondensgaard, Dino Di Carlo, Jennifer Watkins-Yoon

## Abstract

Antibody discovery technologies, essential for research and therapeutic applications, have evolved significantly since the development of hybridoma technology. Various in vitro (display) and in vivo (animal immunization and B cell-sequencing) workflows have led to the discovery of antibodies against diverse antigens. Despite this success, standard display and B-cell sequencing-based technologies are limited to targets that can be produced in a soluble form. This limitation inhibits the screening of function-inducing antibodies, which require the target to be expressed in cells to monitor function or signaling, and antibodies targeting proteins that maintain their physiological structure only when expressed on cell membranes, such as G-protein coupled receptors (GPCRs). A high-throughput two-cell screening workflow, which localizes an antibody-secreting cell (ASC) and a cell expressing the target protein in a microenvironment, can overcome these challenges. To make function-first plasma cell-based antibody discovery accessible and scalable, we developed hydrogel Nanovials that can capture single plasma cells, target-expressing cells, and plasma cell secretions (antibodies). The detection and isolation of Nanovials harboring the antigen-specific plasma cells are then carried out using a flow cytometry cell sorter - an instrument that is available in most academic centers and biopharmaceutical companies. The antibody discovery workflow employing Nanovials was first validated in the context of two different cell membrane-associated antigens produced in recombinant form. We analyzed over 40,000 plasma cells over two campaigns and were able to identify a diversity of binders that i) exhibited high affinity (picomolar) binding, ii) targeted multiple non-overlapping epitopes and iii) demonstrated high developability scores. A campaign using the two-cell assay targeting the immune checkpoint membrane protein PD-1 yielded cell binders with similar EC50s to clinically used Pembrolizumab and Nivolumab. The highest selectivity for binders was observed for sorted events corresponding with the highest signal bound to target cells on Nanovials. Overall, Nanovials can provide a strong foundation for function-first antibody discovery, yielding direct cell binding information and quantitative data on prioritization of hits with flexibility for additional functional readouts in the future.

## Introduction

Since the development of hybridoma technology in the 1970s^1^, antibody discovery methods have significantly evolved. This pioneering technique enabled the production of monoclonal antibodies by merging B cells with myeloma cells, resulting in highly specific and consistent antibodies^2^. Hybridoma technology laid the foundation for many advancements in antibody research and therapeutic applications.

Subsequently, in vivo approaches such as animal immunization and B cell-based technologies were developed^2^. These methods involve immunizing animals to elicit a strong immune response, followed by isolating and sequencing B cells that produce the desired antibodies. Simultaneously, in vitro display technologies, including phage display, yeast display, and ribosome display, emerged as powerful tools for antibody discovery^3^. These techniques utilize immunized or naive antibody libraries displayed on the surfaces of microorganisms, eukaryotic cells, or ribosomes, enabling the rapid screening of large numbers of antibodies against target antigens. These high-throughput methods have significantly accelerated the pace of antibody discovery.

However, significant challenges persist in targeting proteins that are difficult to produce in soluble form. This limitation hinders the discovery of antibodies that require cellular expression of the target for monitoring function or signaling pathways. Moreover, discovering antibodies that interact with membrane-bound proteins, such as G protein-coupled receptors (GPCRs) and ion channels, remains difficult due to their need to be expressed on cells to maintain their physiological structure^4^. GPCRs, in particular, are crucial in various physiological processes and serve as key targets for therapeutic interventions.

To overcome these challenges, high-throughput two-cell screening workflows have been proposed. This method involves compartmentalizing an antibody-producing cell and a target protein-expressing cell within a controlled microenvironment, allowing for the direct screening of antibodies that can bind to cell membrane-associated proteins and/or modulate the function of cell surface targets, thereby enabling function-first antibody discovery. Although some microfluidic and droplet technologies can facilitate two-cell screening workflows, they often suffer from limitations such as low throughput and the necessity for expensive specialized equipment ^5–7^.

We use Nanovials, hydrogel microparticles with cavities that hold nanoliter volumes^8–10^, to sort out plasma cells (PCs) based on secreted IgG that binds to recombinant antigen or antigen expressed on co-localized reporter cell membranes (Figure 1). Studies have shown that differentiation into plasma cells is limited to B cells with high-affinity BCRs^11,12^, reflecting a selection process to optimize humoral immune responses, which are primarily mediated by antibodies where affinity is a crucial factor. Unlike memory B cells, PCs secrete high quantities of immunoglobulins, allowing for cell-based screening assays that enable the prioritization of sequences based on functional information during the initial screening step of antibody discovery workflows. A unique affinity-selected repertoire and soluble secreted antibodies highlight the advantages of PCs as a valuable source of antibody variable region genes.

**Figure 1.**
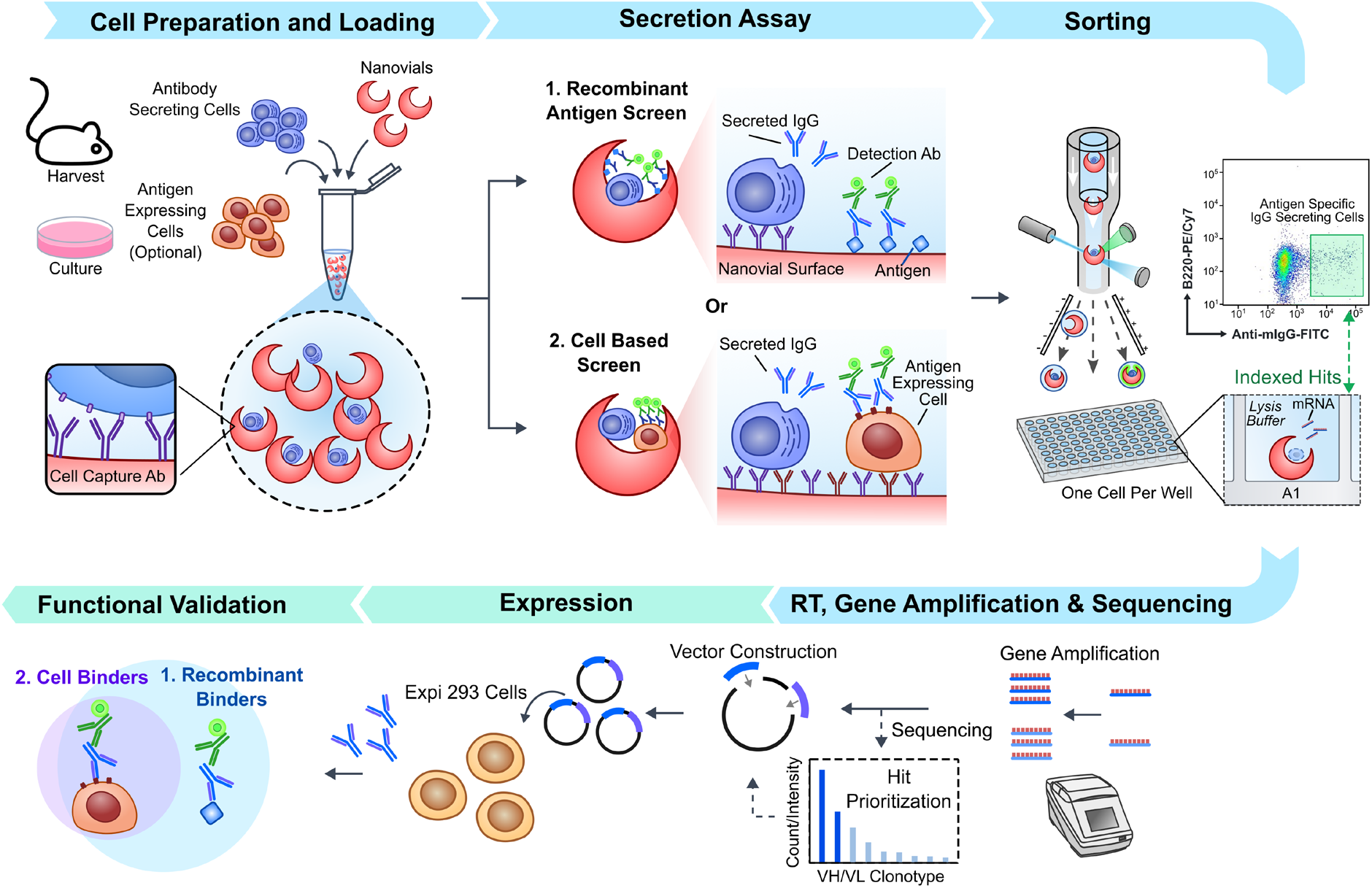
Workflow for plasma cell screening based on antigen-specific IgG secretion. Plasma cells are harvested from bone marrow and loaded into Nanovials, which are pre-modified with capture antibodies. Optionally, Nanovials may also be pre-modified with recombinant antigen. After incubation to allow accumulation of secreted antibodies, antigen-specific antibodies are captured either by bound antigen or by co-loaded cells expressing the antigen, and are subsequently labeled with fluorescent detection antibodies. Antigen-specific ASCs are then individually sorted using flow cytometry. Following sorting, RT-PCR is used to amplify the antibody sequences, which are then sequenced. Initial data analysis helps prioritize clones for re-expression. Selected antibody sequences are incorporated into vectors for transient expression in CHO Expi293 cells, followed by downstream functional validation of the recombinant antibodies.

By functionalizing Nanovials with anti-CD138 antibody to capture plasma cells, antigens to capture secreted IgG, and staining with fluorescent anti-IgG secondary antibody, we isolate single plasma cells, perform single-cell sequencing, and obtain matched heavy and light chain sequences. Importantly, target PCs within Nanovials are sorted using standard fluorescence-activated cell sorters, enabling significantly higher throughput and allowing for the screening of hundreds of thousands of cells per experiment. Unlike other techniques, this method does not require new equipment, thereby increasing accessibility.

Using the Nanovial workflow we first performed two full antibody discovery campaigns to discover novel antibodies against surface-expressed antigens. The process for selecting antigen-specific PCs from cell collection to sorting was complete in one day. We successfully discovered a number of new antibodies, including picomolar binders and several nanomolar binders. Signals present on nanovials containing PCs correlated strongly with the corresponding affinity of re-expressed mAbs from those events. Epitope mapping revealed discovered antibodies targeted different epitopes on the antigen with significant diversity. Because secreted antibodies were used in screening, the majority of sequences discovered had high scores for developability. Cell assays also confirmed that many hits bound native antigen presented in the cell membrane. However, other hits did not bind to cell membrane-presented antigen, motivating us to develop a two-cell assay where both ASCs and antigen-expressing cells were co-loaded in Nanovials. Using the two-cell assay, hybridoma secreting mAbs that bind cell-membrane antigens were substantially enriched in a background of non-specific hybridoma, and new mAbs that target PD-1 with EC50s similar to Pembrolizumab and Nivolumab were identified from PCs. Overall, our results confirm the ability to discover novel high affinity antibodies from PCs using Nanovials, determining functional cell-binding information in the initial screening step and screening at orders of magnitude larger scales and with rapid turnaround time. Nanovial-based workflows improve antibody discovery processes and speed to obtain high affinity, highly functional and developable antibodies.

## Results

We first developed a workflow to specifically capture plasma cells within Nanovials along with any antigen-specific antibodies they secrete (Figure 1). We also developed complementary workflows in which both membrane antigen-expressing cells and ASCs could be co-loaded into Nanovials to sort cells based on antibodies that bound membrane antigen targets (Figure 1). Two types of Nanovials were employed in the assays, biotinylated Nanovials (product # BT-35-A, Partillion Bioscience Corporation) and EZM™ Nanovials (product #s EZ-35-A and EZ-50-A, Partillion Bioscience Corporation). The former type of Nanovials are manufactured to contain biotin molecules on the outer surface and the cavity, while the EZM™ Nanovials have biotin localized solely in the cavity and a larger mean cavity diameter for the same 35 μm outer diameter (20.7 μm vs. 19.1 μm, Figure S1A,C). 50 μm outer diameter EZM™ Nanovials were also used for co-loading two cell types, benefiting from the larger cavity diameter of 31.5 μm (Figure S1B). Nanovials were labeled with streptavidin to enable the coating of biotinylated assay components that in turn permit the capture of ASCs and their secretions. Quality control of Nanovials comprised evaluation of Nanovial size metrics and their uniformity as well as the level of biotin functionalization and biotin uniformity through binding to streptavidin (Figure S1). Fluorophore-conjugated streptavidin coating on Nanovials is confirmed to lead to an increase in fluorescence signal and be uniform (CV <35%) to qualify a batch for use. Multiple permutations of the assay components were evaluated to identify the most-suitable reagents and formats. For cell capture, we found that the use of CD138-specific antibody allowed efficient capture of plasma cells that were isolated from mouse bone marrows, while for hybridoma and target cells anti-CD45 or anti-CD19 antibodies were used to achieve cell capture.

To enable secretion of antibodies by the captured cells, the cells bound to Nanovials were cultured at physiological temperature. The culture time was optimized to minimize cross-talk (contamination of a Nanovial with secretion from a plasma cell on a neighboring Nanovial). Nanovials were then stained with a secondary antibody against mouse IgG, analyzed and sorted using Fluorescence-activated Cell Sorting (FACS) to isolate Nanovials harboring antigen-specific plasma cells as single cells. The secretion capture and detection was attempted in two formats – i) capture using a coated antigen and detection with a fluorescently-labeled anti-mouse IgG; ii) capture with mouse IgG-specific antibody, followed by detection using a fluorescently-labeled antigen. The antigen-on-nanovial format demonstrated the best signal to noise ratio and was employed for the work presented here. This format also more closely parallels the two-cell assay workflow in which target cells expressing membrane antigens are also loaded on Nanovials, and binding on the target cells is detected with fluorescently-labeled anti-mouse IgG. Because Nanovials are larger in size than typical cells we adjusted the drop delay parameter to optimize sorting recovery. Adjusting the drop delay by ‘+0.5’ from the Accudrop setting yielded the highest recovery for the FACSAria III system, where approximately 82% of Nanovials were successfully sorted in the well. We could conduct single-cell RT-PCR to recover heavy and light chain antibody variable genes from cells associated on Nanovials, indicating that the hydrogel Nanovials do not impair sequence recovery.

The above-described Nanovial workflow was then applied to two campaigns aimed at discovering antibodies against cell surface receptors – Ag1 and Ag2. ATX-Gx^TM^ mice harboring human antibody variable genes were immunized with the extracellular domains (ECM) of Ag1 and Ag2 using a repetitive immunizations multiple sites (RIMMS) protocol. The humoral response to the immunization was confirmed by measuring antigen-specific titers in the sera (Figure 2A). Given that most of the germinal center-derived PCs have bone marrow-tropism^13^, bone marrows were harvested for plasma cell isolation. Plasma cells amounted to approximately 0.1% of bone marrow-derived cells in both the campaigns. Using an antigen-on-Nanovial format, plasma cells secreting antigen-specific IgG were identified and sorted as single cells (Figure 2B), processing 22,417 plasma cell events for Ag1 and 23,552 events for Ag2. For Ag1 and Ag2, the antigen-specific signal (cell-loaded Nanovials exhibiting anti-IgG FITC staining within the defined gate) was 3.52% (Figure 2B) and 2.43%, respectively. Incubation on ice instead of a culture step at 37°C to elicit IgG secretion reduced the antigen-specific signal (Figure S2). 130 antigen-specific plasma cells were sorted as single cells in the case of Ag1 and 168 cells were sorted for Ag2 based on fluorescence intensity thresholds defined by the samples incubated on ice (Figure 3A, Figure S2).

**Figure 2.**
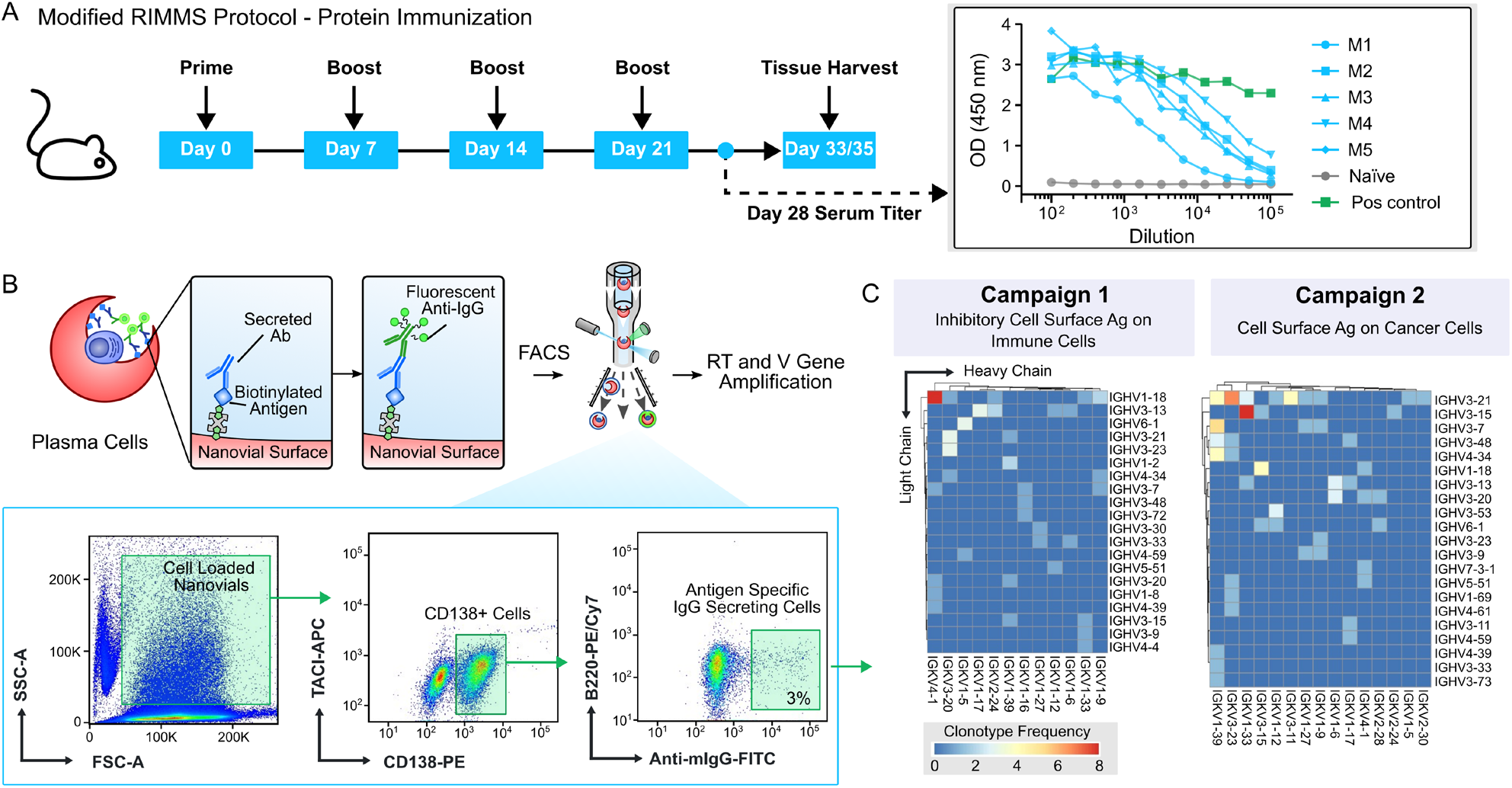
Overview of antibody discovery campaigns using recombinant antigen screening format. **(A)** Overview of immunization protocol and measured serum titers for campaigns 1 and 2. **(B)** Gating strategy used to isolate Nanovials containing CD138+ cells that also exhibit antigen-specific IgG signals. **(C)** Heat maps depicting the distribution of heavy (IGHV-) and light chain (IGKV-) sequences identified post-sort following single-cell sequencing.

**Figure 3.**
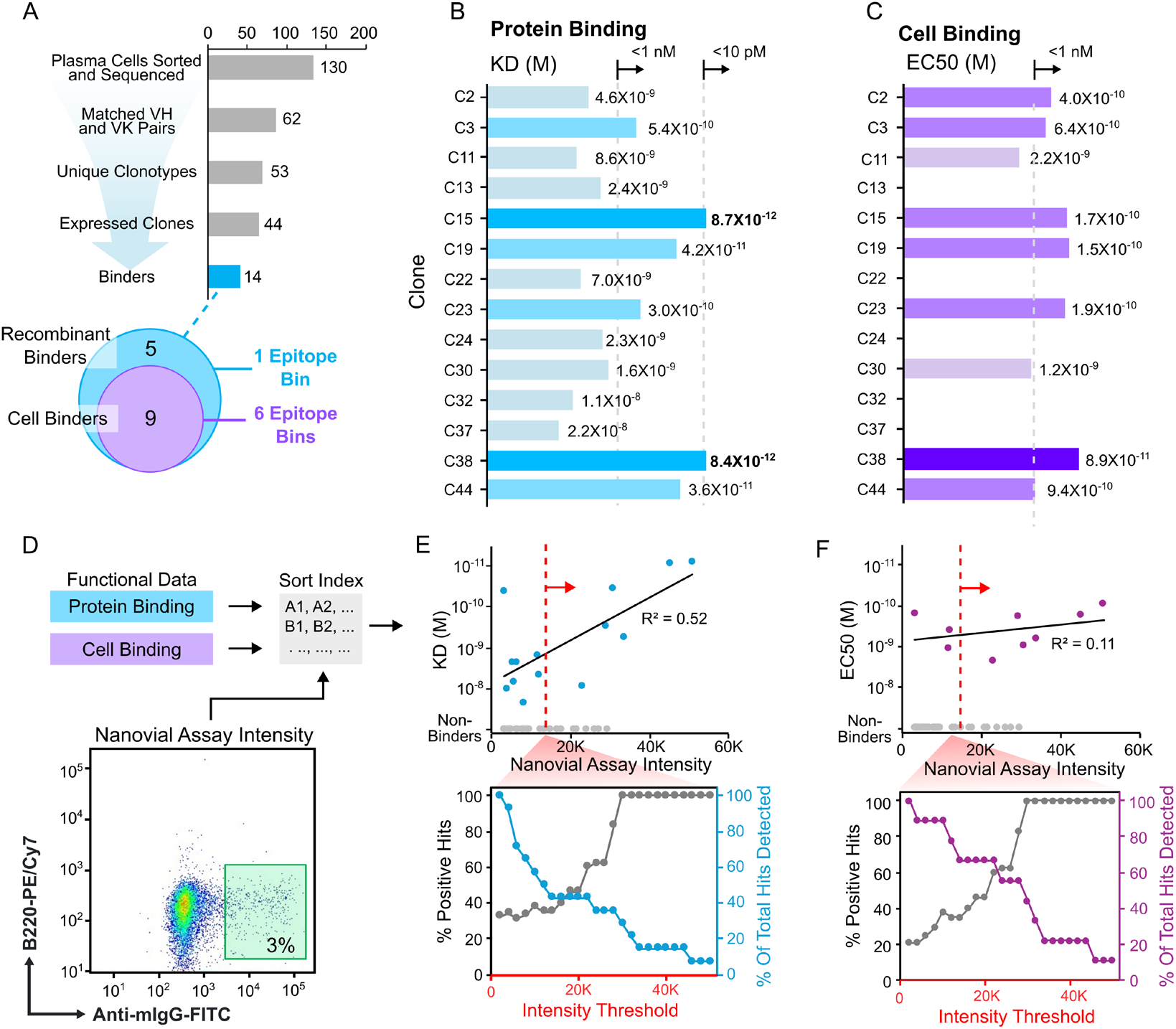
Characterization of antibodies discovered from immunized animals using the recombinant antigen based Nanovial workflow from campaign 1 shown in figure 2. **(A)** Overview of hit attrition post-sort, with a breakdown of identified binders by type and unique epitope bins. **(B)** Dissociation constant (KD) measurements for re-expressed antibodies binding to recombinant antigens, with stronger binders shown in darker blue. **(C)** Cell binding as determined by half-maximum effective concentration (EC50) measurements for re-expressed antibodies. **(D)** Schematic showing the linkage of fluorescence intensity data and functional data using the sorting index. **(E)** Plot of KD versus Nanovial Assay Intensity for all sorted samples (Top). Correlation coefficient (R^2^) for confirmed binders is displayed. The bottom plot shows the fraction of positive hits and the fraction of on-target hits as a function of intensity thresholds. **(F)** Similar to (E), but with comparisons made against EC50 values.

RT-PCR, followed by Sanger sequencing recovered 62 and 102 paired heavy and light chain sequences (with identified V/J genes) in the Ag1 and Ag2 campaigns, respectively (Figure 3A). Importantly, the Ag1 antibody sequences represented 53 clonotypes (20 VH families and 12 VK families) and the Ag2 antibody sequences represented 73 clonotypes (21 VH families and 15 VK families) (Figure 2C, Figure 3A). We observed significant diversity in terms of both V region usage and CDRH3 sequences. Although certain clonotypes were mined in multiples, no single clonotype was dominant.

Across the two campaigns we found a diversity of antibodies with affinity to recombinant antigen and antigen expressed on cells. Antibody affinity to recombinant Ag1 ranged from 22 nM to 8.7 pM (Figure 3B, Figure S2) including 4 sub-nanomolar binders and 2 picomolar binders. Binding to recombinant antigen was also reflected by binding to antigen on cell membranes for both picomolar binders and several other antibodies (Figure 3C, Figure S2), although a fraction of the antibodies did not bind to antigen on cells. The Ag2 campaign also resulted in a number of cell binders (Figure S3). While we initially selected a single threshold to sort putative binders in the Nanovial assay, FACS allowed linking of fluorescence signals of each event with downstream sort locations in the multi-well plates and subsequent functional characterization of the recovered antibodies (Figure 3D). Notably, we found a correlation (R^2^=0.52) between the secretion-associated fluorescence signal on Nanovials (FITC intensity) and affinity to recombinant protein of the re-expressed antibodies corresponding to the sorted events (Figure 3E). Using this data enabled us to retrospectively test out different intensity thresholds for the sort (Figure 3E-F). By picking a threshold intensity of 30,000 instead of the lower sort threshold we would have identified only binders and all binders would have sub-nanomolar affinity. Developability scores for the antibodies were high (Table S1). Antibodies that were identified bound multiple different epitopes of the antigen (Figure 3A, Figure S4) showing a broad repertoire diversity in the plasma cell compartment.

In order to improve the yield of functional antibodies against cell surface membrane-expressed targets we developed a workflow to screen only ASCs that secrete antibodies that bind to target-expressing cells co-loaded on the same Nanovial. Both 35 μm and 50 μm outer diameter Nanovials were able to co-load target Jurkat cells and hybridoma cells producing anti-CD3 antibodies (Figure 4A-B, Figure S5). For both Nanovial types by first loading ASCs and then overloading with the target-expressing cells we were able to maintain high levels of events that contained one ASC and at least one target cell. By flow cytometry we could confirm these events by labeling each cell type with a different dye or fluorescently-labeled antibody (Figure 4A-B). When the double positive events were sorted and imaged, we observed highly enriched populations of Nanovials containing the two cell types (Figure 4B). We validated the two-cell workflow by mixing populations of two hybridoma cell lines (OKT3 and a non-target ovalbumin(OVA)-specific hybridoma) and characterizing binding of secreted antibodies to target CD3-expressing Jurkat cells (Figure 4C). We found selective increase in IgG signal bound to Jurkat cells corresponding to Nanovials also harboring OKT3 cells which was substantially reduced in OVA hybridoma+Jurkat+ Nanovials (Figure 4C,D). When sorted double-positive events gated on high IgG secretion signal were analyzed using microscopy, we observed clear staining on target Jurkat cells reflecting IgG bound to the cell membrane (Figure 4D). By gating on a threshold of IgG intensity (intensity threshold, Figure 4C-v) we enriched OKT3 events from the initial 1:9 spiking ratio to 91.3% of gated events (Figure 4C). Further tuning of the intensity threshold can result in even further increases in purity of the target OKT3 events with a trade-off of reduced percent of the total OKT3 hits being detected. Similar results were observed with anti-CD45 producing hybridoma lines mixed with non-specific hybridoma in 35 μm Nanovials (Figure S5), supporting the generalizability of the assay.

**Figure 4.**
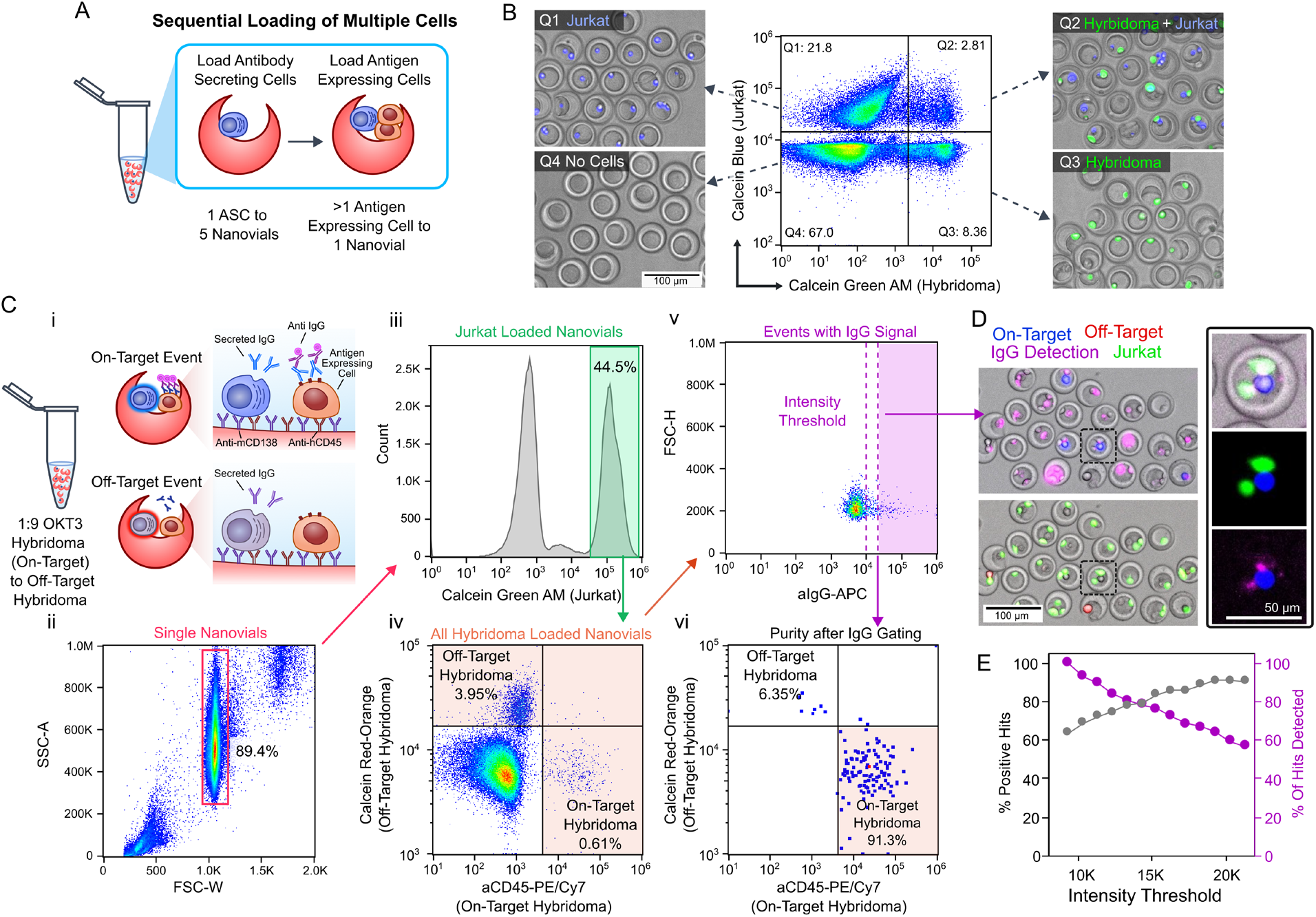
Overview and validation of two-cell Nanovial screening workflow. **(A-B)** Co-loading of Jurkat cells expressing CD3 and OKT3 hybridoma cells producing anti-CD3 antibodies in Nanovials, yielding events with one ASC and at least one target cell, confirmed by flow cytometry and fluorescence microscopy. **(C-D)** Design and validation of the two-cell workflow using hybridoma cell lines secreting both on- and off-target antibodies mixed at a 1:9 ratio. **(C) (i)** Schematic of the assay design and gating strategy to identify co-loaded events and assess purity of events selected based on high IgG signal. On- and off-target hybridoma were labeled with unique dyes to assess percent purity. **(ii)** Forward scatter width (FSC-W) and side scatter area (SSC-A) plots with single nanovials gated. **(iii)** Gating of Nanovial events with Jurkat cells using a calcein AM positive gate. **(iv)** Distribution of hybridoma events on Jurkat-loaded Nanovials showing on-target PE/Cy7 high and off-target calcein red high events. **(v)** Gating and intensity threshold definition for the anti-IgG-APC channel representing labeling of on-target secreted IgG. **(vi)** Based on sorting IgG positive events on-target OKT3 cells are enriched. **(D)** Co-loaded events with high IgG signal were sorted and verified to contain a high percentage of OKT-3 hybridoma and jurkat cells with IgG signal localized on the Jurkat cell surface. **(E)** Plot depicting the effect of an increasing IgG intensity threshold on the purity (% positive hits) and recovery of the on-target hybridoma population. Percent of hits detected (right y axis, pink) is defined as the fraction of cells recovered relative to the most lenient gating threshold.

After developing the two-cell assay with known hybridoma lines, we applied it to screen plasma cells from mice immunized with human PD-1, an immune checkpoint control on T cells (Figure 5A). Human Raji cells with forced overexpression of PD-1 were employed as target cells. We sorted Nanovial events that contained (i) Raji cells, (ii) plasma cells from immunized animal bone marrows and (iii) high fluorescence signal associated with captured IgG on target cells (Figure 5B). Following sequencing and re-expression of hits this campaign resulted in four cell binding clones with EC50s of less than 1 nM, similar to EC50s of 0.30 nM and 0.35 nM for the clinically-used checkpoint inhibitor antibodies, Pembrolizumab and Nivolumab (Figure 5C). As for our recombinant antigen assay we found the intensity of anti-IgG signal observed by flow cytometry was a good predictor of the binding propensity of discovered antibodies (Figure 5D).

**Figure 5.**
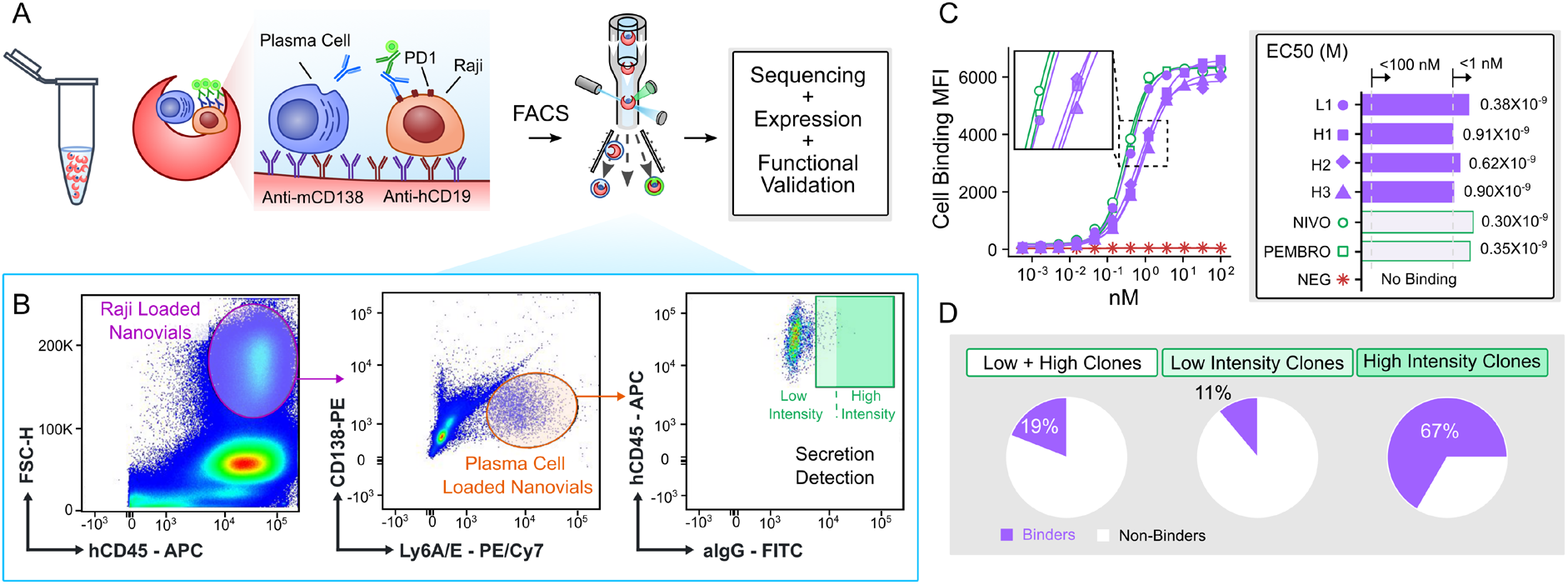
Antibody discovery campaign for PD-1 using the two-cell Nanovial assay. **(A)** Schematic of the Nanovial format with anti-mouse CD138 and anti-human CD19 for the screen of plasma cells secreting antibodies that bind PD-1 on the cell membrane of Raji cells. High IgG events are sorted by FACS, re-expressed and functionally validated. **(B)** Gating strategy for Raji-loaded Nanovials containing plasma cells and anti-IgG signal. A threshold for the fluorescence intensity for an event was used to sort and classify the function of recovered clones. (**C**) Cell binding curves of four cell binding antibodies discovered with EC50 values shown in comparison with Pembrolizumab (PEMBRO) and Nivolumab (NIVO). H1-H3 are events from the High Intensity gate while L1 is from the Low Intensity gate. NEG is the negative control non-PD-1 binding antibody. **(D)** Pie charts showing the percentage of clones that were cell binders for different intensity gates, indicating a higher intensity threshold yielded a larger fraction of events that were true cell binders.

## Discussion

Nanovials enable function-first based screening of ASCs and simultaneous discovery and ranking of novel antibody sequences based on properties such as affinity or cell binding. The compatibility with standard reagents and flow cytometry instruments makes these approaches accessible and rapid, operating at sorting throughputs of up to 500 events per second, which is sufficient to sort through hundreds of thousands of ASCs in a single day. The flexibility of the Nanovial platform, which allows for adjustments in cell capture antibodies and nanovial cavity size, can support the development of new two-cell workflows. This adaptability is particularly useful for screening antibodies that target other cell membrane antigens, such as G-protein coupled receptors (GPCRs), which are among the most significant classes of druggable targets.

The sensitive and quantitative signals from the flow cytometry readout also provide unique information that is helpful for prioritizing sequences for further investigation. For example, fluorescence signal strength on Nanovials reflected the affinity of the secreted antibodies to recombinant antigen, even though this signal might be expected to report on a combination of secreted quantity (and local antibody concentration) and antibody affinity. The capability to link this information to particular sorted clones, using index sorting, provides a guide for further follow up of hits, and can be used to retroactively create new gating thresholds for sequencing depending on the number of sequences that are desired to be re-expressed. Increased fluorescence intensity on target-expressing cells bound within Nanovials also was an indicator of true cell binding of an antibody in the two-cell assay format, substantially increasing the fraction of clones that were functional cell binders upon re-expression. Antibody binding information can be maintained and linked to sequences in other downstream sequencing pipelines, such as single-cell RNA-seq using the 10x Chromium, by using the secretion-encoded single-cell sequencing (SEC-seq) technique ^14–16^. In SEC-seq the bound secreted IgG in a Nanovial is labeled with an oligonucleotide barcode which enables linkage of the quantity of bound IgG with the V(D)J sequences for the same corresponding cell. Further refinement in affinity-based measurements on the Nanovial are also possible by capturing secreted antibody and separately measuring the amount of secreted antibody and antigen bound to each Nanovial – normalizing for secretion differences.

Two-cell assay workflows are particularly challenging using current approaches (Table S2) and Nanovial-based workflows open up many new possibilities for developing different two-cell assay types. Our initial demonstration focused on binding assays, however, assays that incorporate reporter cells that may respond to antibody binding, or antibody internalization assays are also possible. Image-activated cell sorting, which is now becoming more widely available^17^, could be enabling for other more sophisticated workflows.

Our results also suggest the benefits of screening plasma cells in antibody discovery campaigns. We see significant heterogeneity in the different clones identified and do not see many repeat clones, indicating access to a diverse antibody pool. The plasma cell compartment may differ from the memory B cell compartment by reflecting predominantly affinity-matured antibodies. This is supported by the many high affinity binders we identified across multiple campaigns, including two picomolar binders to Ag1. The secreted antibodies, if they aggregate or otherwise lack stability would not create strong signals on Nanovials and therefore are not selected for in our screen, ensuring some baseline quality for developability which can be lacking from other approaches like antigen baiting of B cells, in vitro display or AI-prediction based approaches (Table S2).

Overall, we anticipate the scale of screening and functional antibody performance data that is acquired at the screening stage of the Nanovial workflow will guide antibody discovery and engineering, and help train new AI predictive models^18,19^. The accessibility of Nanovials can level the playing field, offering a platform that fosters further innovation in antibody discovery. This advancement has the potential to lead to the development of improved therapeutic molecules reaching clinical application.

## Methods

### Immunizations

To generate antibody against Ag1 and Ag2, two cohorts comprising each five ATX-GK mice were immunized. ATX-GK mice are transgenic animals that can produce antibodies with human variable regions. Cohorts 1 and 2 were immunized subcutaneously with a weekly dose of His-tag-carrying Ag1 and Ag2, respectively. After a total of 5 injections for cohorts 1 and 2, mice were sacrificed, and bone marrows harvested for all the animals (Figure 2A).

For titer measurement, mouse sera were prepared and used in an ELISA against Ag1 or Ag2 proteins. Briefly, for each cohort, Ag1 or Ag2 proteins were coated on an ELISA plate. Plates were blocked with 3% BSA buffer for 1hr and after PBS wash, sera dilutions were transferred to the ELISA plate and incubated for 1hr. After PBS wash, a secondary antibody specific for mouse IgG labeled with horse radish peroxidase (HRP) was added to the ELISA plate for 1 hr. After PBS wash, 3,3’,5,5’-Tetramethylbenzidine (TMB) solution was added, and reaction was then stopped using sulfuric acid.

### Bone marrow processing and CD138+ cell isolation

To process bone marrow, the mouse femur and tibia (F/T) were cleaned in a sterile environment to remove epithelial and muscle tissue and then placed in sterile and cold RPMI. A mortar sterilized with 70% ethanol was placed in the hood, and the F/T were transferred into it. The F/T were sterilized by incubation in cold 70% ethanol for about 30 seconds. The ethanol was then aspirated and discarded, and the F/T were washed with cold RPMI to remove any traces of ethanol. The RPMI was aspirated and discarded, and 15 mL of R10+ (RPMI + 10% FBS + 1:1000 BLI’s DNA clean-up) was added per 5 sets of F/T. Using a pestle sterilized with 70% ethanol, the F/T bones were broken to release the bone marrow (BM), which was transferred with the R10+ into a 50 mL tube through a 70 μm cell strainer. An additional 10 mL of R10+ was added to the broken bones, mixed gently to release any residual BM, and transferred into a second 50 mL tube through a 70 μm cell strainer. This step was repeated 2-4 times until the bones turned white, with the resulting cell suspension equally distributed into the first and second 50 mL tubes. The BM cell suspension was incubated for 10 minutes at room temperature, centrifuged for 6 minutes at 400g, and the supernatant was discarded. To lyse red blood cells, 5 mL of cold ACK RBC lysis buffer was added and incubated on ice for 30-45 seconds. Subsequently, 30 mL of cold MACS buffer was added to each tube and centrifuged at 400g for 6 minutes. The supernatant was discarded, and 10-15 mL of cold MACS buffer was added. The solution was filtered through a 70 μm cell strainer, and the filter was washed to recover any remaining cells. The cell count was determined, and the protocol proceeded to CD138+ plasma cell isolation.

To isolate CD138+ plasma cells, the Miltenyi Biotec CD138+ Plasma Cell Isolation Kit (Miltenyi Biotec CAT# 130-092-530) was followed with some modifications. All centrifuge steps were performed at 350g for 7 minutes at 4°C, two rounds of positive selection using MS columns were employed and 100 μL was subtracted from the calculated volume to account for residual pellet volume for all volumes of buffer. CD138+ cell isolation proceeded in steps according to the kit instructions, first depleting non-plasma cells and then enriching CD138+ cells through two separate magnetic columns. Ten microliters of collected cells were set aside for counting, and the remaining cells were prepared for further protocols such as staining. After isolation, the cells were centrifuged for 6 minutes at 400g, the supernatant was discarded, and the cell pellet was resuspended in the required buffer or medium to proceed to the screening workflow.

### Nanovial preparation and functionalization

Two types of Nanovials were used for the workflow. For the initial antigen-binding campaigns with Ag1 and Ag2, product # BT-35-A (Partillion Bioscience Corporation) was used. For two-cell assay workflows EZM™ Nanovials (product #s EZ-35-A and EZ-50-A, Partillion Bioscience Corporation) were used. For Campaigns 1 and 2, Nanovials were prepared as follows: Nanovials were pipetted from a stock concentration of 6.2 × 10^6^ Nanovials/mL into an Eppendorf tube and diluted 2X with wash buffer (0.5% BSA, 1% Penicillin-Streptomycin, 0.05% Pluronic F127 in PBS) containing 0.5 μg/mL APC-Cy7-labeled streptavidin (BD cat# 554063) and 59.5 μg/mL of unlabeled streptavidin (ThermoFisher Scientific cat# S888). The mixture was incubated at room temperature for 60 minutes. The Nanovials were washed three times by filling the tube with wash buffer and centrifuging at 300 g for 3 minutes at room temperature. The pelleted Nanovials were then resuspended in wash buffer to reach the stock density and diluted 2X with additional wash buffer containing 24 μg/mL of anti-mCD138 IgG-Biotin and 12 μg/mL of biotinylated antigen. This suspension was incubated at room temperature for 2.5 hours. Finally, the Nanovials were washed three times with wash buffer.

For the two-cell assays, Nanovials were prepared as follows: Nanovials were pipetted from a stock concentration of 4 × 10^6^ Nanovials/mL into an Eppendorf tube and rinsed with wash buffer. The tube was topped off with 1 mL of wash buffer and centrifuged at 200 g for 5 minutes at room temperature. The supernatant was aspirated and Nanovials were suspended to a concentration of 1 × 10^6^ Nanovials/100 μL. An equivalent volume of wash buffer containing 400 μg/mL of unlabeled streptavidin (Thermofisher CAT#S888) was added and the mixture was incubated at room temperature for 90 minutes on a rotator set at the lowest speed. The Nanovials were washed twice by topping off the tube with wash buffer and centrifuging at 200 g for 5 minutes at room temperature. The pelleted Nanovials were then incubated in 500 μL of wash buffer containing 15 μg/mL of anti-mCD138 IgG-Biotin– targeting mouse plasma cells and 15 μg/mL of anti-hCD45 IgG-Biotin – targeting human Jurkat cells. This suspension was incubated at room temperature for 2.5 hours on a rotator set at the lowest speed. Finally, the Nanovials were washed twice with wash buffer, by centrifuging at 200 g for 5 minutes at room temperature, and aspirating the supernatant, to prepare them for cell loading.

### Cell loading on Nanovials

Loading nanovials was carried out at 4°C in a refrigerator. The pelleted nanovials were combined with 100 μL of suspended plasma cells per 1×10^6^ nanovials. The appropriate number of cells was added to create a 1: 10 (one-cell assays) or 1:20 (two-cell assays) ratio of cells to nanovials. This mixture was incubated at 4°C for 60 minutes with gentle mixing once after 30 minutes. The cell-loaded nanovials were pelleted by centrifugation at 200g for 5 minutes at 4°C, and the supernatant was aspirated. For two-cell assays, subsequently, 20-50-fold excess of reporter cells were added to the pelleted nanovials, following the same gentle mixing procedure. The mixture was incubated again at 4°C for 60 minutes with gentle mixing once after 30 minutes. The cell-loaded nanovials were then pelleted by centrifugation at 200g for 5 minutes at 4°C, and the supernatant was aspirated. Finally, the pellet was resuspended in R10 medium (RPMI + 10% FBS) for secretion incubation as described below.

### Secretion incubation and staining

The following antibodies were used for cell staining/blocking prior to the secretion incubation (20-30 minutes at 4°C in 2% FBS/PBS): i) one-cell discovery campaigns – Fc-block (5 μg/ml; BD Biosciences cat# 553142) and anti-mouse TACI-APC (3 μg/ml; STEMCELL Technologies cat# 60116AZ); ii) two-cell hybridoma experiment – anti-mouse CD138-PE-Cy7 (BioLegend cat# 142514), anti-mouse CD138-PE (BioLegend cat# 142504) and anti-mouse IgG2a/2b (10 μg/ml; BCR blocking); iii) two-cell PD-1 campaign-Fc-block (5 μg/ml; BD Biosciences cat# 553142), anti-mouse Ly6A/E-PE/Cy7 (5 μg/ml; BioLegend cat# 108113), anti-mouse IgG1 (20 ug/ml; BCR blocking) and ant-mouse IgG2a/2b (20 ug/ml; BCR blocking). Following incubation, the samples were washed twice with 2% FBS/PBS, centrifuged at 200-300g for 3-5 mins at 4°C, and the supernatant was aspirated. For the secretion incubation, cell-loaded nanovial pellet was resuspended in 1 mL (one-cell assays) or 10 mL (two-cell assays) of R10 medium per 1×10^6^ nanovials. A no-secretion control sample and a positive control sample was aliquoted and stored on ice or at 4°C. The tubes were incubated for 60 or 90 mins for one-cell or two-cell assays, respectively, at 37°C with 5% CO_2_ on a slant. After incubation, the tubes were removed from the incubator, or refrigerator for control conditions, and centrifuged at 200-300 g for 3-5 minutes at 4°C. The supernatant was aspirated, and the pellets were resuspended in sort buffer (2% FBS, 1% Penicillin-Streptomycin, 0.05% Pluronic F127 in PBS) containing the following antibodies i) one-cell discovery campaigns – anti-mouse IgG1-FITC (5 μg/ml; anti-mouse IgG2a/b-FITC (5 g/ml) and anti-mouse CD138-PE (4 ug/ml; BioLegend cat# 142504); ii) two-cell hybridoma experiment-anti-mouse IgG1-FITC (5 ug/ml), anti-mouse IgG2a/b-FITC (5 ug/ml) and and anti-human CD45-APC (Invitrogen cat# MA5-17688); iii) two-cell PD-1 campaign -anti-mouse IgG1-FITC (5 μg/ml), anti-mouse IgG2a/b-FITC (5 μg/ml) anti-mouse CD138-PE (5 μg/ml; BioLegend cat# 142504) and anti-human CD45-APC (Invitrogen cat# MA5-17688). The mixture was incubated at 4°C for 30 minutes. Following incubation, the samples were washed twice with sort buffer, centrifuged at 200-300g for 3-5 mins at 4°C, and the supernatant was aspirated. The pellets were then resuspended in the appropriate volume of sort buffer to achieve a flow/sort sample with an event rate equal to or lower than 1,000 events per second.

### Two-cell secretion assay

For the OKT3 hybridoma two-cell secretion assay, 50 μm Nanovials were used. Biotinylated Nanovials were incubated with a final concentration of 200 μg/mL streptavidin for 30 minutes while rotating and then washed. Nanovials were then modified with 20 μg/mL biotin anti-mouse CD138 volume and 20 μg/mL biotin anti-human CD45 for hybridoma and Jurkat loading, respectively.

After washes, the nanovials were loaded with a OKT3 spiked hybridoma mixture that was 10% OKT3 and 90% OVA hybridoma. Both cell types were stained separately before being combined and loaded into nanovials. The OKT3 hybridomas were double-stained with Calcein Blue and PE-Cy7 anti-mouse CD45, while the OVA hybridomas were stained with Calcein Red-Orange. Both hybridomas were also blocked with anti-mouse IgG2a during the staining step. The hybridoma mixture was loaded in a 1:5 hybridoma to Nanovial ratio and incubated on ice for 1 hour to allow hybridomas to bind to Nanovials via anti-mouse CD138 antibody. Calcein Green, AM labelled Jurkats were subsequently added to the hybridoma-loaded samples in a 5:1 Jurkat to Nanovial ratio and incubated on ice for 30 minutes to allow Jurkats to bind to nanovials via anti-human CD45 antibody. Samples were then run through a 20 μm strainer to remove unbound cells and reverse-strained to recover Nanovials. The samples were then incubated in a 37°C, 5% CO_2_ incubator for 30 minutes to accumulate secretions. During this time, the positive control was incubated with concentrated OKT3 hybridoma conditioned media. After washes, the samples were stained with PE anti-mouse IgG2a and washed a final time.

### Nanovial screening assay and flow cytometry

#### FACSAriaTM

All flow cytometry sorting for plasma cell campaigns (both antigen binding and two-cell workflows) and 4B2 hybridoma studies were performed using a FACSAria™ Fusion or FACSAria™ III (BD Biosciences) equipped with a 100 μm nozzle.

##### Sample Preparation and Compensation

Nanovial samples were diluted to approximately 1e6 Nanovials/mL in Partillion Sort Buffer for analysis and sorting. Voltages for different fluorescence channels were set using cell-loaded nanovial samples, and compensation was carried out using negative controls (blank compensation beads; ThermoFisher Scientific cat# 01-2222-42) and positive controls (compensation beads stained using fluorescently-labeled antibodies).

##### Drop Delay and Calibration

Drop delay value for Nanovials was optimized using the following protocol: The drop-delay on the instrument was set using Accudrop beads (BD Biosciences cat# 345249), and a plate alignment for sorting into a 96-well plate was performed. A 96-well plate was loaded with 100-200 μL of wash buffer in each well for sorting Nanovials. 5,000 Nanovials were sorted into each well by manually adjusting the drop delay value in eight increments and eight decrements of 0.25 from the drop-delay value obtained using Accudrop beads. A known number of counting beads (ThermoFisher Scientific cat# C36950) were added to each well, and the contents were transferred into tubes for flow cytometry analysis. Each tube of samples was run on the FACS instrument, and at least 500 Nanovials were acquired. The data was analyzed to determine the number of counting beads and Nanovials in each sample. Since the number of added counting beads was known, the total number of Nanovials sorted into each well was back-calculated. The efficiency of sorting at each drop-delay value was then calculated using the formula: (sorted events/expected events) * 100.

##### Gating Strategy

The following gating strategy was used to identify antibody secreting cells of interest: One-cell workflow: (1) cell-loaded nanovial population based on high forward scatter area and side scatter area, (2) exclusion of any free cells based on nanovial stain, (3) inclusion of only plasma cell-loaded nanovials based on plasma cell markers, (4) IgG signal.

Two-cell workflow (hybridomas): (1) reporter cell-loaded Nanovial population based on high forward scatter area and antigen presenting cell marker signal (2) Hybridoma stain positive population, (3) IgG signal on antigen presenting cells

Two-cell workflow (plasma cells): (1) reporter cell-loaded Nanovial population based on high forward scatter area and antigen presenting cell marker signal, (2) inclusion of only plasma cell-loaded Nanovials based on plasma cell markers, (3) IgG signal.

#### SONY MA900

Instruments and Configuration. OKT-3 hybridoma two-cell studies were performed using a SONY MA900 cell sorter equipped with a 130-micron sorting chip (SONY Biotechnology) and 4 lasers (violet, blue, green, red).

##### Laser and Filter Setup

Blue, green, and red lasers were turned on for OKT-3 hybridoma two-cell studies. Data was obtained using the Standard Filter set on the MA900 instrument. Default gain settings were used (40%) except for the following: FSC 1, BSC 26.5%, FL1: 33%, FL2 41%, FL5 48%, FL10 42.5%. The threshold channel was FSC set at 0.5%

##### Sample Preparation and Compensation

Nanovial samples were diluted to 0.5e6 Nanovials/mL in Partillion Wash Buffer for analysis and sorting. Samples were compensated using negative controls (unstained cells) and positive single-color controls of OKT3 cells labeled with the following fluorophores: Calcein-AM (Thermo Fisher), Calcein red-orange (Thermo Fisher), anti-CD45 APC (Biolegend), anti-CD45 PE-Cy7 (Biolegend)

##### Drop Delay and Calibration

Drop delay was first configured using the automated calibration process. For 50 μm Nanovials, the sort delay settings were manually offset to achieve the highest purity and recovery. The sort delay was adjusted in increments of ±1 and evaluated by sorting 50 Nanovials into a 96-well plate and manually counting them. An offset of -2.0 with the single-cell, 3-drop sort mode provided the best performance for 50 μm Nanovials.

##### Sample Pressure

A sample pressure setting of 4 was used for all analysis and sorting processes.

##### Gating Strategy

The following gating strategy was used to identify antibody secreting cells with antigen presenting cells on Nanovials: (1) Singlet Nanovials gated based on FSC-W, (2) antigen presenting cell signal positive population, (3) Hybridoma stain positive population, (4) IgG signal on antigen presenting cells.

##### Data Acquisition and Analysis

Data acquisition was performed using the appropriate software for the respective flow cytometers, and all analyses were conducted using FlowJo software (FlowJo LLC). Statistical analysis and data visualization were performed using GraphPad Prism (GraphPad Software).

### Sequencing and re-expression of mAbs

cDNA synthesis and VH/VL gene amplification were conducted according to previously published protocols^20^. RNA from single plasma cells within the sorted nanovials was reverse transcribed using the SuperScript™ IV First-Strand Synthesis Kit (Thermo Fisher Scientific cat# 18091200). Custom primers designed for ATX-GK mice were utilized to amplify human *Igh* and *Igk* V gene transcripts through two rounds of nested PCR. The PCR products were analyzed on 2% agarose gels, purified, and sequenced via Sanger Sequencing by a commercial sequencing facility. Following the identification of the V gene families, family-specific custom primers were used to re-amplify the PCR1 product, which was then cloned into independent expression vectors for human IgG1 heavy chain and kappa light chain. Sequence-verified constructs were transiently transfected into Expi293™ cells (Life Technologies) to produce the antibodies, which were subsequently purified from the culture supernatants using protein A-Sepharose.

### Assays of mAb function

Various assays for mAb function were conducted on the re-expressed antibodies as outlined below.

#### SEC HPLC

Size exclusion chromatography (SEC) was performed with a YMC Diol-200 8 x 300 mm column (Cat.no # DL20S05-3008WT) on an Agilent 1200 series HPLC instrument. The running buffer was 20mM sodium phosphate, 400mM NaCl pH 7.0 at a flow rate of 0.3 mL/min. For freeze-thaw stability; samples frozen at -80C for 20 minutes, then thawed at room temperature for approximately 20 minutes.

#### CE SDS

Capillary electrophoresis sodium dodecyl sulfate (CE-SDS) was performed on a LabChip GX II instrument using Protein Express 200 (Perkin Elmer, #760499) and Protein Express Assay Reagent Kit (Perkin Elmer, # CLS960008). The reagents and chip were prepared according to the manufacturer’s instruction. Briefly, the reducing sample buffer was prepared by mixing 1 M dichlorodiphenyltrichloroethane with Protein Express Sample Buffer, while the nonreducing buffer consisted of only Protein Express Sample Buffer. Samples were mixed with reducing or non-reducing buffer and denatured at 80 °C for 10 min. Samples were centrifuged at 2,000 g for 1min to remove air bubbles before placing in the LabChip GXII instrument for analysis.

#### BLI for binding affinity

Kinetic experiments were performed on Carterra LSA with HBST buffer (10mM HEPES pH7.4, 150mM NaCl, 3mM EDTA, 0.05% Tween 20) as the running buffer. Capture sensor-chip was prepared by covalently immobilizing goat anti-human IgG capture antibodies (Jackson ImmunoResearch) onto a HC30M chip using a standard amine-coupling procedure. Briefly, chip was activated with 33 mM s-NHS and 133 mM EDC in 100 mM MES pH 5.5 for 7 minutes. Antibodies at 10 mg/ml in acetic acid buffer pH 4.5 were used for printing for 10 min. The printed chip was then quenched with 1 M ethanolamine pH 8.5 for 7 min. For kinetics analysis, purified recombinant his tagged human or cyno PD-1 (ACRO) at a concentration from 0.025 nM to 500 nM (a serial 3-fold dilution) was injected sequentially. For each concentration, there was 5 Min association followed by 10 Min dissociation. Results were processed and analyzed in Carterra LSA Kinetics Software. The kinetic data was referenced with the interstitial reference spots and double-referenced to a buffer cycle, and then fit globally to a 1:1 binding model to determine their apparent association and dissociation kinetic rate constants (ka and kd values). The ratio kd/ka was used to derive the KD value of each antigen/mAb interaction, i.e. KD=kd/ka.

#### Epitope binning

High-throughput epitope binning was performed using real-time label-free biosensors (Carterra LSA) to sort large panels of mAbs into bins based on their ability to block one another for binding to the antigen. In a pairwise epitope binning analysis, Ag1 and antibody 2 (analyte antibody) are sequentially applied to the sensor chip (HC200M) covalently pre-loaded with antibody 1 (ligand antibody). An increase in response upon exposure to the analyte antibody indicates non-competition between the two antibodies, whereas a lack of change in the signal indicates competition. Antibodies having the same blocking profiles towards others in the test set are grouped into one bin.

Community network plots are used to explore clustering of mAbs that share similar but not necessarily identical competition profiles. Rather than relying strictly on the sandwiching/blocking assignments in the heat map, as the Bin network plots do, hierarchical clustering is applied to the sorted heat map to generate dendrograms, which progressively group mAbs.

#### DSF

Thermal stability of mAbs was assessed via nano differential scanning fluorimetry (nanoDSF) on Prometheus Panta. Each sample was measured in duplicate. Melting temperatures of the antibodies were detected during heating in a linear thermal ramp (0.5 °C/min, 25-95 °C). Data was analyzed using the Panta Analysis software. The unfolding transition points were determined from changes in the emission wavelengths of tryptophan fluorescence at 350 and 330 nm.

#### AC-SINS

Gold nanoparticles (Ted Pella, 15705-20) were washed with water. Antibody mixture of 80/20 (v/v) capture antibody/non-capture antibodies (Jackson Immuno Research Labs) was buffer exchanges into 20 mM Sodium Acetate pH 4.5 to a concentration of 500 μg/ml. To prepare 1 ml coated particles, 900 μL of gold nanoparticles were incubated overnight with 100 μL of antibody mixture for 90 min at RT. After antibody coating, thiolated PEG (MW: 2000 Da) were used to quench the beads. The beads were then concentrated 10 fold in PBS. 10 μL 10x concentrated particles solution was incubated with 100 μL of 40 μg/mL of antibody samples on a 384-well polypropylene plate for 2 hrs RT. The plate was then quickly spun down at 3000 rpm and scanned from 510 to 580 nm nm in increments of 2 nm on Synergy Neo2 Multi-Mode Plate Reader (BioTek). Values reported are averages of duplicate wells and are sample red shift wavelengths at maximum absorbance subtracting the blank reference (PBS only). Greater red shifts indicate increased self-interaction.

#### BVP ELISA

The method was similar as reported by Hötzel et al. (2012). Briefly, baculovirus particles (BVP, Lake Pharma) was diluted 1:100 in 50 mM sodium bicarbonate (pH 9.5). After overnight incubation of 50 μL of BVP on ELISA plates (3369; Corning) at 4 °C overnight, unbound BVPs were aspirated from the wells. All remaining steps were performed at RT. The plate was blocked with 100 μL of blocking buffer (PBS with 1% BSA) for 1 h before three washes with 100 μL of PBS. Next, 50 μL of 2.5 ug/mL testing antibodies was added to the wells and incubated for 1 h followed by washes with 100 μL of PBS. HRP-conjugated goat anti-human IgG antibody at 1:1000 (Jackson ImmunoResearch) was used as the secondary antibody, and incubated for 1 h followed by washes as before. Finally, 100 μL of TMB substrate (34021; Fisher Scientific) was added to each well and incubated for 6 min. The reactions were stopped by adding 50 μL of 2 M sulfuric acid to each well. The absorbance was read at 450 nm and BVP score determined by normalizing absorbance by control wells with no test antibody. Fold-over background was determined using an average of buffer-only wells. All measurements were performed in triplicate.

#### Cell binding assay

Cells expressing Ag1 and Ag2 were tested in binding assays with anti-Ag1 and anti-Ag2 antibodies respectively at concentrations from 100 nM to 0.6 pM (a serial 3-fold dilution) for 45 min on ice. Cells were then incubated with the secondary antibody R-Phycoerythrin AffiniPure Goat Anti-Human IgG (Jackson Immunoresearch 109-115-098). The data was acquired on FACSCanto II. Median fluorescence intensities (MFI) were plotted against the concentrations of the antibodies. EC50 was derived from fitting to 4 parameter dose-response curve.

## Acknowledgements

The authors acknowledge funding from the National Institutes of Health (Grant #R43GM144000 and R44GM144000, PI J.D.). The authors would like to thank Alex Wu, Wei-Ying Kuo, Lucy Liu, Lisa Sherman, Dalton Markrush, Erica Teng, Cédric R. Weber, Enrique Batitay, Stefanie Chan, Caity Goshert, Sarah Beatty, Ellie Knecht, and Alan Nhan for their assistance with conducting experiments and providing materials and data.

