## Supplemental Information for "Function-first discovery of high affinity monoclonal antibodies using Nanovial-based plasma B cell screening"

1. Alloy Therapeutics
2. Partillion Bioscience Corporation
3. University of California, Los Angeles

**Figure S1. Quality control (QC) results and a summary table of key QC metrics.**

**Figure S2. Example flow plots showing antigen-specific IgG signal on Nanovials without secretion incubation and with secretion incubation.**

**Figure S3. Cell binding of re-expressed antibodies to Ag2.**

**Figure S4. Properties of re-expressed antibodies to Ag1**.

**Figure S5. Two-cell Nanovial workflow and results with anti-CD45 producing hybridoma.**

**Table S1: Developability scores for antibodies discovered from plasma cell secretion campaign #1.**

**Table S2: Comparison of antibody discovery techniques.**

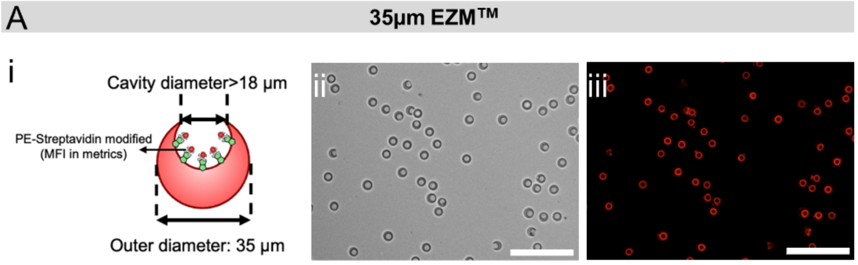


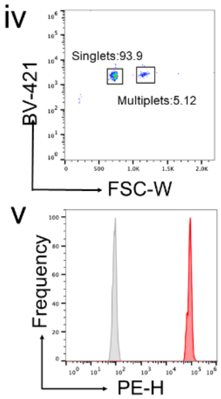


| Key QC metrics | Size (um) | CV | Cavity Size (µm) | MFI | MFI CV | Multiplet % |
|---|---|---|---|---|---|---|
| This batch | 34.7 | 3.2 | 20.7 | 98549 | 22.6 | 5.12 |
| Specification (35µm) | 35±5 | <10% | >18 | >10000 | <35% | <10% |

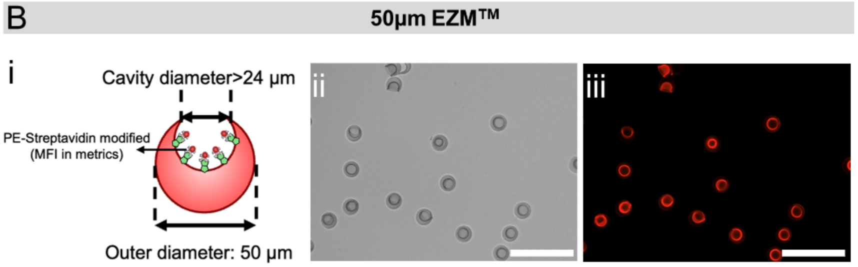


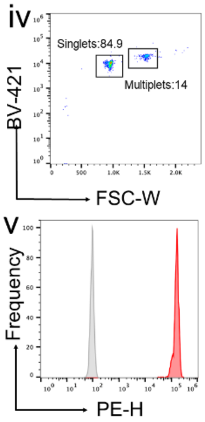


| Key QC metrics | Size (um) | CV | Cavity Size (µm) | MFI | MFI CV | Multiplet % |
|---|---|---|---|---|---|---|
| This batch | 51.8 | 4.3 | 31.5 | 143301 | 13.6 | 14 |
| Specification (35µm) | 50±5 | <10% | >24 | >30000 | <35% | <20% |

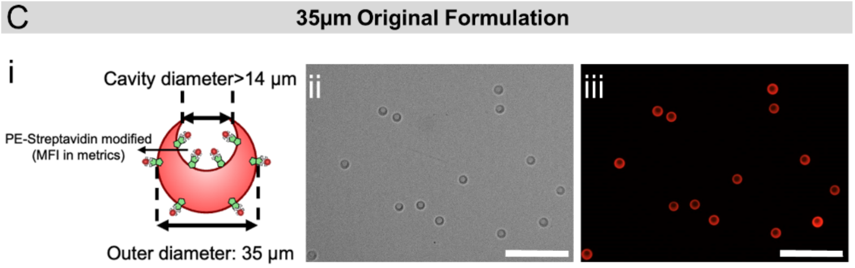


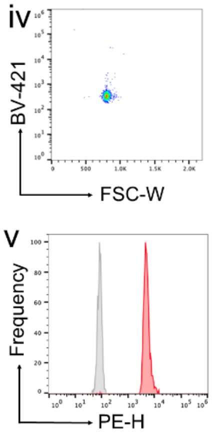


| Key QC metrics | Size (um) | CV | Cavity Size (µm) | MFI | MFI CV |
|---|---|---|---|---|---|
| This batch | 33.8 | 4.3 | 19.1 | 5864 | 25.7 |
| Specification (35µm) | 35±5 | <10% | >14 | >2000 | <35% |

**Figure S1. Quality control (QC) results and a summary table of key QC metrics.** A) Metrics for 35µm EZM™ Nanovials, B) 50 µm EZM™ Nanovials and C) 35 µm biotinylated (original formulation) Nanovials. For each panel, the subfigures are as follows: **i)** A schematic representation of the of the Nanovials including definitions of the key QC metrics. **ii)** A brightfield image of the Nanovials. **iii)** Fluorescence images for the same particles in subfigure ii). **iv)** A scatter plot showing the properties of the Nanovials on an FSC-W versus BV-421 plot, used to identify the singlet/multiplet ratio. and **v)** A PE-H histogram of PE-streptavidin stained (red peak) and unstained Nanovials (grey peak), quantifying the available binding sites on the Nanovials. All scale bars are 200 µm.

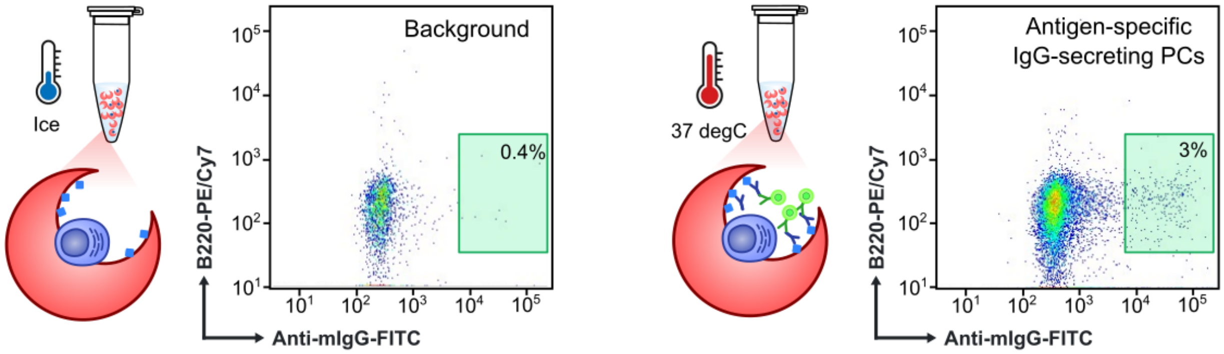


**Figure S2. Example flow plots showing antigen-specific IgG signal on Nanovials without secretion incubation (Left) and with secretion incubation (Right).** Schematics show plasma cells on Nanovials with antigen coating the Nanovial surface. With incubation at 37°C secreted IgG binding to antigen is shown, followed by anti-IgG labeling. Flow plots show anti-IgG signal on the x axis and cell stain (B220 PE/Cy7) on the y axis. Nanovials with plasma cells incubated on ice led to a reduced fraction of antigen-specific IgG events compared to experimental samples.

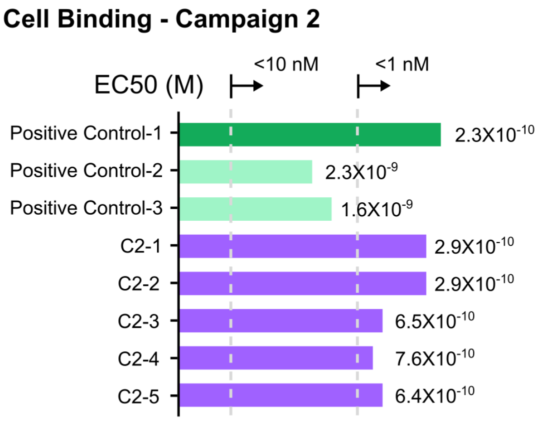


**Figure S3. Cell binding of re-expressed antibodies to Ag2.** EC50 of antibodies discovered in the campaign against Ag2 (C2-1 through C2-5) compared with three positive control antibodies. Discovered clones all had EC50 values less than 1nM.

A

| # | KD | EC50 | Type | Community |
|---|---|---|---|---|
| POS-1 | | | Positive Control | 1 |
| C11 | 8.6E-09 | 2.2E-09 | Sample | 2 |
| C15 | 8.7E-12 | 1.7E-10 | Sample | 2 |
| C19 | 4.2E-11 | 1.5E-10 | Sample | 2 |
| C23 | 3.0E-10 | 1.9E-10 | Sample | 2 |
| POS-2 | | | Positive Control | 2 |
| C2 | 4.6E-09 | 4.0E-10 | Sample | 3 |
| C22 | 7.0E-09 | 0.0E+00 | Sample | 3 |
| C44 | 3.6E-11 | 9.4E-10 | Sample | 3 |
| C3 | 5.5E-10 | 6.4E-10 | Sample | 4 |
| C13 | 2.4E-09 | 0.0E+00 | Sample | 5 |
| C24 | 2.3E-09 | 0.0E+00 | Sample | 6 |
| C30 | 1.6E-09 | 1.2E-09 | Sample | 6 |
| C32 | 1.1E-08 | NB | Sample | 7 |
| C37 | 2.2E-08 | NB | Sample | 7 |
| C38 | 8.4E-12 | 8.9E-11 | Sample | 8 |

B

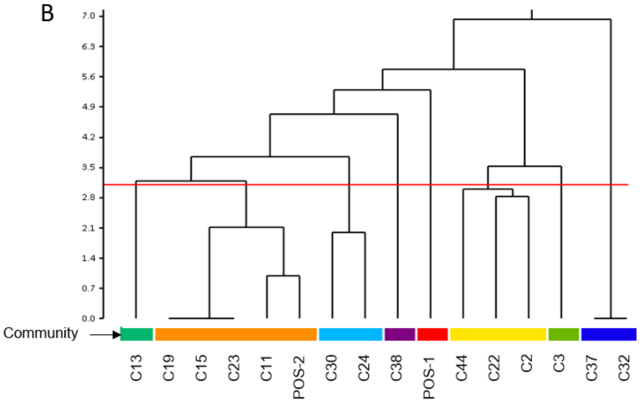


**Figure S4. Properties of re-expressed antibodies to Ag1**. **(A)** Binding affinity to recombinant antigen (KD) and cell binding (EC50) values for discovered clones clustered by community bins. Two known positive controls were included for reference. **(B)** Bins and dendrogram showing relationships between antibody binding epitopes showing a diversity of function.

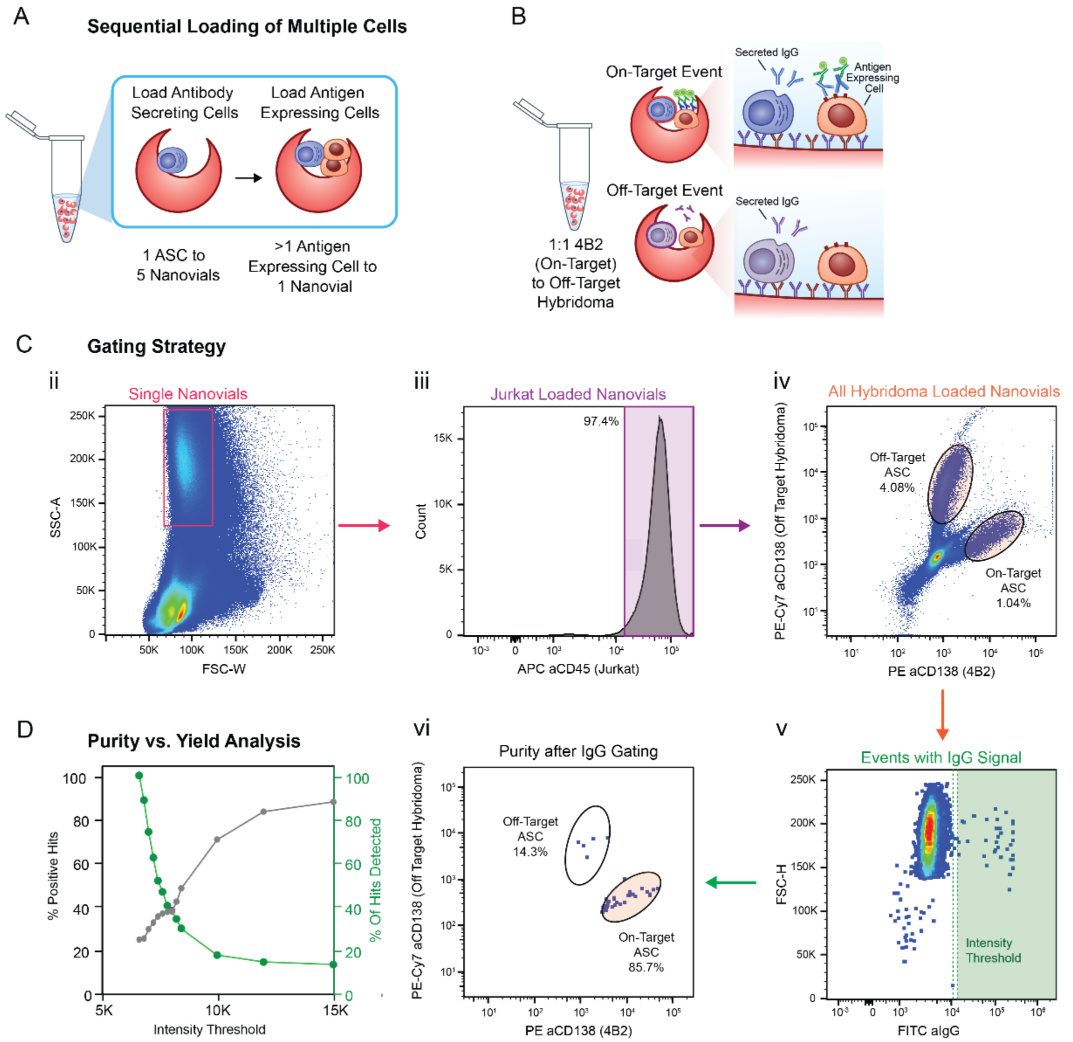


**Figure S5. Two-cell Nanovial workflow and results with anti-CD45 producing hybridoma. (A)** Schematic of the loading process for the two-cell assay. **(B)** Schematic showing specific secretion and detection on target antigen-expressing cells (Jurkat cells) in Nanovials. 4B2 are anti-CD45 secreting hybridoma. (**C**) Gating strategy showing gating of single 35 µm Nanovials (ii) loaded with Jurkat cells (iii) and then either of the ASCs (4B2 and off-target hybridoma, iv). Gating on IgG secretion signal (v) is used above the intensity threshold to determine the on-target and off target populations that are enriched (vi). (**D**) With increasing intensity thresholds an increasing percentage of positive 4B2 cells would be enriched, exceeding 80%, with a tradeoff of reduced number of hits that would be detected.

**Table S1:** Developability scores for antibodies discovered from plasma cell secretion campaign #1.

| Sample Info | | KD (M) | Cell Binding | Thermal Stability (nDSF) | | | Developability | | | | | | | |
|---|---|---|---|---|---|---|---|---|---|---|---|---|---|---|
| | | | | nDSF (C) | | Turbidity (C) | Normalized Score | Salt Conc. (M) | Normalized Score | LSA polyreactivity (binding score) | | | | |
| # | Community Bin | hu Antigen | EC50 (nM) | Tm1 | Tm2 | IP | AC-SINS | HIC | BVP ELISA | HSP90 | HEK-CP | HEK-IMP | KLH | LPS |
| 2 | 3 | 4.61E-09 | 4.0E-10 | 69.09 | 74.72 | 73.32 | 0.02 | 0.87 | 0.06 | 0.00 | 0.00 | 0.00 | 0.00 | 0.00 |
| 3 | 4 | 5.46E-10 | 6.4E-10 | 67.79 | 73.83 | 73.21 | -0.01 | 0.64 | 0.16 | 0.00 | 0.00 | 0.00 | 0.00 | 0.00 |
| 11 | 2 | 8.60E-09 | 2.2E-09 | 70.51 | 77.40 | 74.14 | 0.04 | 0.72 | 0.08 | 0.00 | 0.00 | 0.00 | 0.00 | 0.00 |
| 13 | 5 | 2.37E-09 | N.P. | 68.65 | 74.49 | 70.74 | 0.32 | 0.40 | 0.07 | 0.00 | 0.25 | 0.25 | 0.25 | 0.00 |
| 15 | 2 | 8.66E-12 | 1.7E-10 | 70.78 | 77.30 | 75.55 | 0.10 | 0.77 | 0.04 | 0.00 | 0.00 | 0.00 | 0.00 | 0.00 |
| 19 | 2 | 4.24E-11 | 1.5E-10 | 70.16 | 76.16 | 73.86 | 0.06 | 0.78 | 0.05 | 0.00 | 0.00 | 0.00 | 0.00 | 0.00 |
| 22 | 3 | 7.02E-09 | N.P. | 70.54 | 77.04 | 79.19 | -0.02 | 0.98 | 0.02 | 0.00 | 0.00 | 0.00 | 0.00 | 0.25 |
| 23 | 2 | 2.95E-10 | 1.9E-10 | 70.06 | 79.29 | 80.00 | 0.13 | 0.86 | 0.06 | 0.00 | 0.00 | 0.00 | 0.00 | 0.00 |
| 24 | 6 | 2.27E-09 | N.P. | 68.83 | 75.37 | 71.56 | 0.18 | 0.34 | 0.08 | 0.00 | 0.50 | 1.00 | 0.50 | 0.00 |
| 30 | 6 | 1.59E-09 | 1.2E-09 | 68.80 | 75.40 | 71.55 | 0.20 | 0.34 | 0.09 | 0.00 | 0.75 | 1.00 | 0.75 | 0.00 |
| 32 | 7 | 1.06E-08 | 0.0E+00 | 70.50 | 77.92 | 80.69 | -0.03 | 0.85 | 0.06 | 0.00 | 0.25 | 0.00 | 0.00 | 0.25 |
| 37 | 7 | 2.18E-08 | 0.0E+00 | 69.18 | 75.20 | 71.71 | -0.03 | 0.71 | 0.02 | 0.00 | 0.00 | 0.00 | 1.00 | 0.00 |
| 38 | 8 | 8.38E-12 | 8.9E-11 | 70.20 | 77.48 | 75.68 | 0.00 | 1.02 | 0.06 | 0.00 | 0.00 | 0.00 | 0.00 | 0.00 |
| 44 | 3 | 3.55E-11 | 9.4E-10 | 65.48 | 71.27 | 67.66 | -0.02 | 0.89 | 0.07 | 0.00 | 0.00 | 0.00 | 0.00 | 0.00 |

**Table S2:** Comparison of antibody discovery techniques.

| Technique | Antigen Baiting[1-2] | Hybridoma[1,3] | In Vitro Display[1] | Microfluidic Screening[1,4-6] | Nanovials (current work) |
|---|---|---|---|---|---|
| **Directly Screen Plasma Cells** | No | No | No | Yes | Yes |
| **Repertoire Diversity** | High[#] | Low | High | High | High |
| **Formats** | Ag Binding | Ag Binding, Two-cell | Ag Binding | Ag Binding, Two-cell | Ag Binding, Two-cell |
| **Affinity Ranking** | Possible | Yes | No | Yes | Yes |
| **Native VH and VK Pairing** | Yes | Yes | No | Yes | Yes |
| **Drug-like Form** | No | Yes | No | Yes | Yes |
| **Native Ag Presentation** | No | Yes | No | Yes | Yes |
| **Workflow Time** | 2-5 Days | 3 Weeks | 3 Weeks | 2-5 Days | 2-5 Days |
| **Readout** | FACS | ELISA | ELISA | Custom Instrument | FACS |
| **Throughput** | >1M | 100-1000's | >1M | 10k-1M[+] | 50k-1M |
| **Accessibility** | High | Med | Med | Low | High |

* Throughput represents the number of events in a typical experiment for one day of screening.

* Workflow time represents the time from cell harvest to sequence recovery.

* Readout. FACS – fluorescence activated cell sorting. ELISA – enzyme linked immunosorbent assay. Custom Instrument refers to specialized instruments with additional capital costs.

* Formats. Ag Binding indicates ability to determine binders to recombinant antigen. Two-cell indicates ability to assay binding to cell membrane proteins, reporter cell function, etc.

* Affinity Ranking. Ability to use optical / fluorescence data to prioritize hits and optimize recovery of hits based on affinity.

*Drug-like form. Indicates the ability to find highly developable candidates. Non-aggregating candidates that are easily produced and stable.

### Excluding the diversity of the affinity-matured plasma cell compartment.

+ Commercially available instruments have demonstrated throughputs
